# Choice-induced biases in perception

**DOI:** 10.1101/043224

**Authors:** Long Luu, Alan A Stocker

## Abstract

Illusions provide a great opportunity to study how perception is affected by both the observer's expectations and the way sensory information is represented^1,2,3,4,5,6^. Recently, Jazayeri and Movshon^7^ reported a new and interesting perceptual illusion, demonstrating that the perceived motion direction of a dynamic random dot stimulus is systematically biased when preceded by a motion discrimination judgment. The authors hypothesized that these biases emerge because the brain predominantly relies on those neurons that are most informative for solving the discrimination task^8^, but then is using the same neural weighting profile for generating the percept. In other words, they argue that these biases are “mistakes” of the brain, resulting from using inappropriate neural read-out weights. While we were able to replicate the illusion for a different visual stimulus (orientation), our new psychophysical data suggest that the above interpretation is likely incorrect: Biases are not caused by a read-out profile optimized for solving the discrimination task but rather by the specific choices subjects make in the discrimination task on any given trial. We formulate this idea as a conditioned Bayesian observer model and show that it can explain the new as well as the original psychophysical data. In this framework, the biases are not caused by mistake but rather by the brain's attempt to remain ‘self-consistent’ in its inference process. Our model establishes a direct connection between the current perceptual illusion and the well-known phenomena of cognitive consistency and dissonance^9,10^.

---

We first tested whether the original illusion generalizes to other stimulus variables. We replicated the main experiment of the original study^7^ using, however, a visual orientation stimulus consisting of a circular array of small line segments (see Fig. 1a). After stimulus presentation subjects first had to indicate whether the overall orientation of the array was clockwise (cw) or counter-clockwise (ccw) of a discrimination boundary (discrimination task), and then had to reproduce their perceived overall orientation of the array by adjusting a reference line (estimation task). We tested three levels of stimulus uncertainty by adjusting the width of the distribution the individual line segments were sampled from.

**Figure 1:**
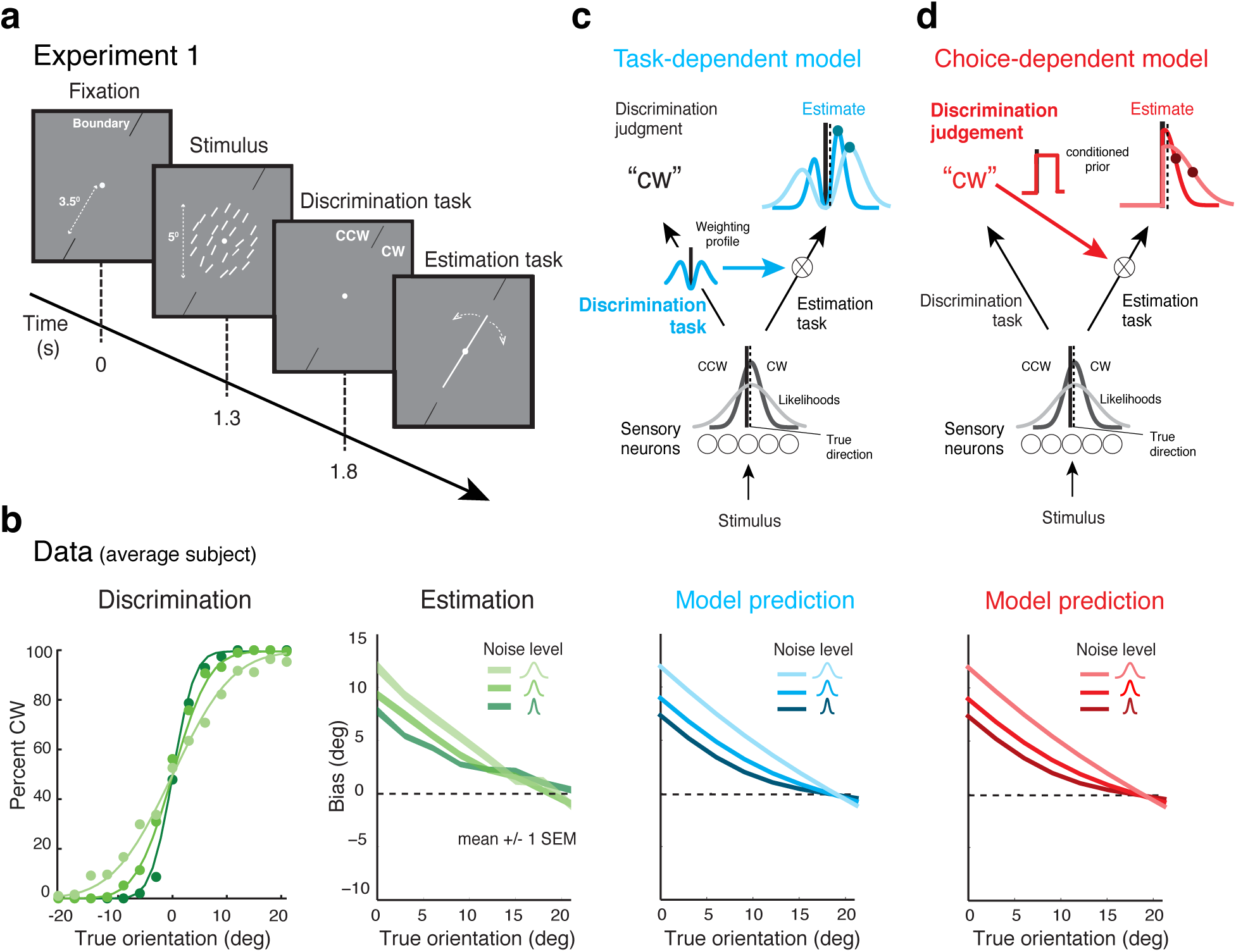
*Generalizaron of the illusion and potential model explanations.* **(a)**Experiment 1: The design was essentially identical to the original study except that we used an orientation stimulus. **(b)** Data for the average subject (N=5; see Extended Data Figure 1 for individual subject data). Subjects’ behavior was very similar compared to the original study, both in terms of the discrimination performance and their estimation biases (compare, *e.g*., to Figs. 2b and 3c in Jazayeri and Movshon^7^). Biases are repulsive, away from the discrimination boundary (only shown for correct trials). **(c)** Task-dependent model by Jazayeri and Movshon^7^. The estimation biases are assumed to arise from a *bimodal sensory weighing profile* aimed at optimally solving the discrimination task. **(d)** Alternative choice-dependent model proposed by Stocker and Simoncelli^11^. In contrast, this model assumes that the biases arise because a subject's *judgment* in the discrimination task (*e.g*. ‘cw’) re-enters the subsequent perceptual estimation process in form of a conditioned prior.

We found that subjects’ response behavior in both the discrimination and the estimation task was very similar to the results of the original study. Discrimination performance monotonically depended on the level of stimulus uncertainty and, more importantly, perceived stimulus orientations showed the same repulsive biases away from the discrimination boundary with larger biases for stimulus orientations closer to the boundary and for higher levels of stimulus uncertainty (Fig. 1b).

According to Jazayeri and Movshon^7^ these biases are the result of a selective readout of the population of neurons representing the sensory evidence. They hypothesized that the brain preferentially weighs signals from those neurons whose responses are most informative with respect to the fine discrimination task, but is then compelled to use the same (bimodal) weighting profile when performing the secondary estimation task. In their model, weighing (*i.e*. multiplying) an unimodal sensory representation centered at the true stimulus orientation with a bimodal weighting profile, and applying a winner-takes-all mechanism to the weighted response leads to stimulus estimates that show repulsive biases (Fig. 1c). Jazayeri and Movshon demonstrate that with a particular choice of the weighting function the model can be fit to the observed experimental biases. A distinct characteristic of the model is that the weighting profile must be established right at the beginning of each trial (as soon as the discrimination boundary is shown and the task is defined, Fig. 1a), and then both the discrimination as well as the estimation outcome simply follow from two *independent* feedforward processes based on the same sensory information and the same weighting profile.

An alternative and, as we will demonstrate, more appropriate explanation of the illusion is to consider the two tasks as causally *dependent,* where a subject's judgment in the discrimination task directly conditions the inference process of the subsequent estimation task (Fig. 1d). More specifically, we assume that subjects trust their choice in the discrimination task such that in the estimation process, they only consider stimulus values that are consistent with this choice. This behavior can be mathematically formulated as a conditioned Bayesian observer model^11^. The model jointly accounts for subjects’ behavior in both the discrimination as well as the estimation task. The model makes the fundamental assumption that along a task sequence based on the same sensory evidence, human perception attempts to remain *self-consistent* by conditioning the subsequent inference process on the subject's preceding decisions.

While both models can account for the original dataset^7,11^ and the results of Experiment 1, they obviously provide fundamentally different explanations for the illusion. We run two additional experiments designed to distinguish the two model hypotheses. Experiment 2 was identical to Experiment 1 except that at the beginning of each trial, subjects were explicitly reminded of the total range within which the stimulus orientation would occur in the trial (gray arc, Fig. 2a). Because the optimal weighting profile for the discrimination task does not depend on the stimulus range (except for very small ranges), the model by Jazayeri and Movshon does not predict any change in shape nor magnitude of the estimation biases compared to the measured biases in Experiment 1. In contrast, the self-consistent Bayesian observer model per definition depends on the stimulus range (*i.e*. the stimulus prior). Thus the model predicts a shift of the crossover point of the estimation bias curves towards the discrimination boundary under the assumption that the explicit representation of the stimulus range provides subjects with a better, and presumably narrower, estimate of the stimulus prior. Indeed, the measured estimation bias curves in Experiment 2 show a clear and consistent shift of the crossover point toward the discrimination boundary (Fig. 2b), much in agreement with the predicted shift by our proposed model, but not the model by Jazayeri and Movshon.

**Figure 2:**
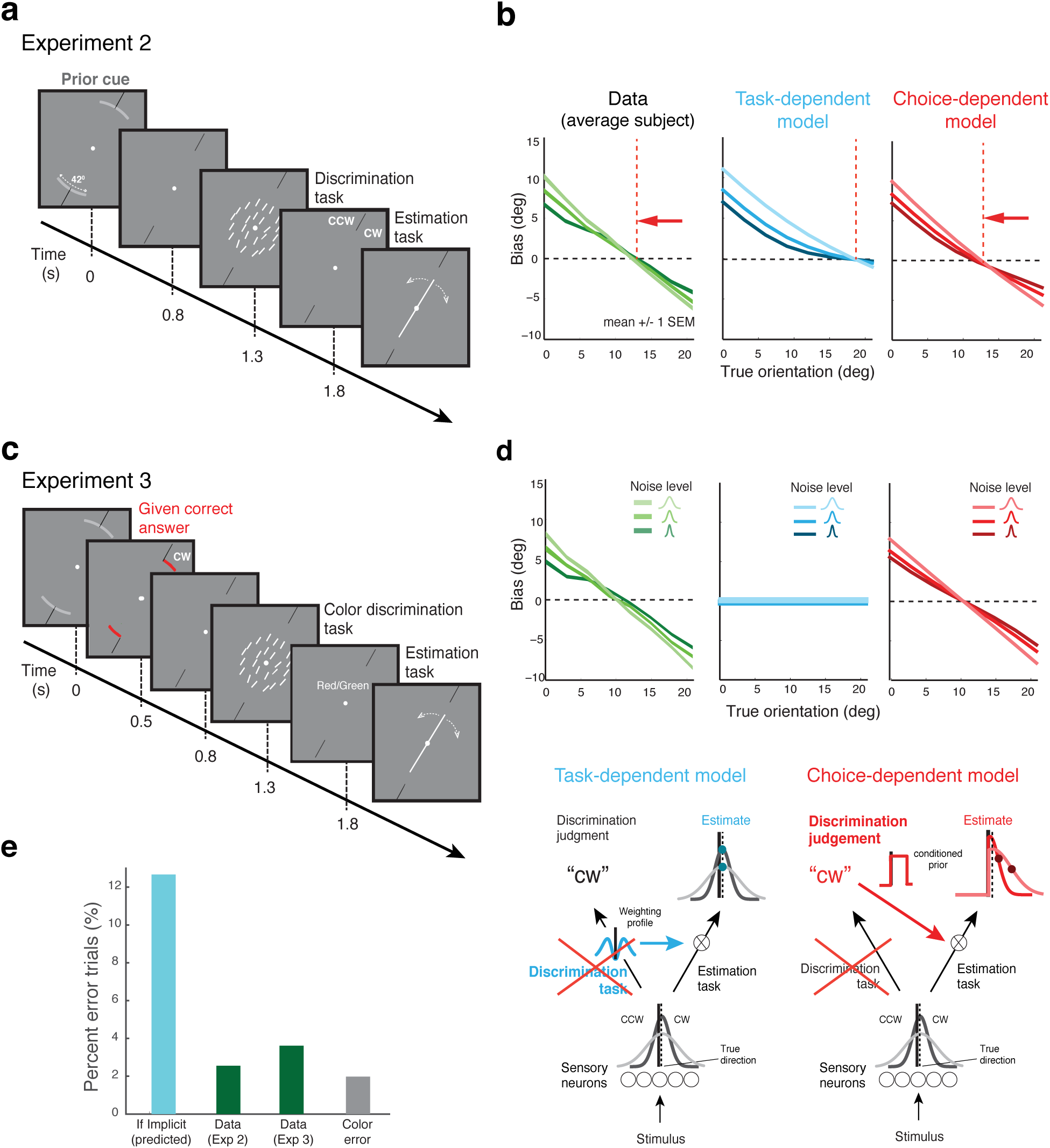
*Probing the two model hypotheses*. **(a)** Experiment 2 was identical to Experiment 1, except that at the beginning of each trial subjects were shown the total range within which the stimulus orientation would occur in the trial (gray arc). The stimulus distribution relative to the discrimination boundary was uniform and the same throughout all three experiments. **(b)** Subjects’ bias curves (average subject, see Extended Data Figure 3 for individual subject data) were shifted towards the discrimination boundary compared to the curves obtained in Experiment 1 (Fig. 1b). Because the optimal weighting profile does not depend on the stimulus range, this change is not predicted by the model of Jazayeri/Movshon. It is, however, correctly accounted for by our proposed conditioned Bayesian observer model assuming that explicitly showing the range leads to a more accurate (and more narrow) estimate of the prior distribution. **(c)** Experiment 3: subjects were provided with the correct answer for the orientation discrimination task *before* the stimulus was presented. Subjects performed an unrelated color discrimination task instead, where they needed to remember the color (red/green) of the cue indicating the correct answer. There was no correlation between the color (red/green) and the given correct answer (’cw’ or ‘ccw’). **(d)** The measured bias curves were very similar to the curves measured in Experiment 2 (average subject; see Extended Data Figure 3 for individual subject data). This is inconsistent with the task-dependent model of Jazayeri/Movshon because without the need to perform the discrimination task, there is no rationale for assuming the same bimodal weighting profile necessary for explaining these biases. In contrast, the self-consistent Bayesian observer model would, by definition, treat a ‘given answer’ identical to a judgment the subject had made him/herself. As a result, the model essentially predicts the same bias curves as those in Experiment 2, which is supported by the data (there is a slight difference in the predictions because the given answer was always correct while a subjective judgment naturally can be incorrect). **(e)** To exclude the scenario in which subjects simply ignored the given answer and implicitly performed the orientation discrimination task themselves instead, we compared the fraction of incongruent trials (trials where the discrimination judgment was inconsistent with the estimate) across Experiments 2 and 3 with the predicted fraction if they were indeed making implicit judgments. The fraction was much lower and of comparable size to subjects’ error rates in the color discrimination task, suggesting that the errors are simply due to erroneous memory retrieval.

With Experiment 3 we directly targeted the key difference between the two models by separating the discrimination judgment from the discrimination task. Subjects were no longer asked to perform the fine orientation discrimination task but instead were directly given the correct answer at the beginning of each trial by a colored cue (Fig. 2c). For control reasons they still had to perform a color discrimination task that, however, was completely unrelated to the orientation stimulus. As Fig. 2d shows, subjects’ estimation biases in Experiment 3 are very similar to the biases in Experiment 2.

This rules out the model by Jazayeri/Movshon because, in order to account for the data, it would need to assume the same weighting profile (optimized for the discrimination task) for Experiments 2 and 3, even though subjects no longer need to perform the discrimination task in Experiment 3. Instead, we expect the model to predict no (or at least very different) estimation biases. In contrast, because the subjects still have access to a discrimination judgment signal, the conditioned Bayesian observer model predicts estimation biases that are essentially identical to those in Experiment 2 (a small difference remains because the given answer is always correct while the subjective judgments naturally can be incorrect). Note that both Experiment 2 and 3 were conducted on the same set of subjects (with half of the group doing Experiment 2 first and the other half starting with Experiment 3). We can also rule out that subjects may have ignored the given answer and implicitly performed the orientation discrimination task instead, and thus still have applied a bimodal weighting profile leading to the observer biases. If this were the case, then the expected rate of inconsistent trials (*i.e*. trials in which the estimated orientation was not in agreement with the given correct answer) would be much larger than what we measured. Instead, the rates for Experiment 2 and 3 are comparable and are of the same magnitude as the error rates for the (irrelevant) color discrimination task, suggesting that these errors simply reflect memory retrieval errors (Fig. 2e).

## Summary and conclusion

We were able to replicate the new perceptual illusion reported by Jazayeri and Movshon^7^ for a different visual stimulus, which suggests that the observed estimation biases may reflect a general perceptual illusion. However, our new data strongly suggest that the original explanation for these biases is incorrect. We find that it is the subjects’ individual judgments rather than the demand for an optimal performance in the discrimination task (as proposed by Jazayeri and Movshon) that causes the biases in the subsequent estimation task. This choice-dependent behavior can be well formalized with a conditioned Bayesian observer model in which a subject's estimate of the stimulus value is conditioned not only on the sensory evidence but also on the subject's judgment in the preceding discrimination task^11^. The across-subject differences in behavior are well described by meaningful variations in subjects individual noise levels and their knowledge of the accurate stimulus prior (Extended Data Figure 1 and 3). This is also in stark contrast to the model by Jazayeri and Movshon where the individual differences in the readout profile lack any clear interpretation. Finally, the model not only accounts for subjects’ average estimates, but also for the full distribution of the estimates both in correct and incorrect trials (see *e.g*. Extended Data Figure 2).

The implications of our results are of significance. First, our data suggest that subjects did not distinguish between a decision outcome they generated themselves (as in Experiment 2) and a decision outcome that was given to them (Experiment 3), implying that a human observer in such a perceptual task sequence treats their own subjective judgment as if it were true. Computationally, this is interesting because it is clearly a non-optimal behavior with regard to perceptual accuracy (obviously, the subjective judgment can be wrong) yet has the fundamental advantage that the subject is “self-consistent” along the sequential inference chain. The proposed self-consistent observer model may indeed reflect a more general framework for understanding perception and behavior within the time-dependent causal structures of natural environments: The brain fuses incoming sensory evidence with both prior expectations and its current internal hypotheses about the structure of the world, and thus, by ensuring self-consistency in drawing conclusions from the same sensory evidence, generates and maintains a robust and consistent interpretation of the external world. Second, our new interpretation of the illusion creates a direct link to the well known phenomena of cognitive consistency^9^ and dissonance^10^. It leads to the interesting and exciting hypothesis that there is a very general and unifying description of human sequential decision making behavior that transcends the perception/cognition boundary. Finally, results from recent physiological recordings show that choice-related signals are fed back along the perceptual processing pathway all the way to early sensory areas^12^’^13^. The new observer model may provide an explanation for the computational purpose and perceptual consequences of these feedback signals: to guarantee that the perceptual process remains self-consistent.

## Acknowledgements

This work was supported by the University of Pennsylvania and the National Science Foundation of the United States of America (CAREER award 1350786).

## Methods

### Experimental setup

Ten subjects with normal or correct-to-normal vision (6 males, 4 females; one non-naïve) participated in the experiments. One subject (male) was excluded from the analysis because he failed to correctly execute the estimation task. All subjects provided informed consent. The experiments were approved by the Institutional Review Board of the University of Pennsylvania under protocol #819634. The number of subjects and trials were chosen such that the statistical power in our data was similar to the one of the original study^7^, which allowed a fair comparison. Details of the experimental setup can be found in the Supplement.

### Conditioned Bayesian observer model

Let *θ* be the true stimulus orientation, *m* the noisy sensory measurement gathered from the stimulus, and *C* = {’cw’, ‘ccw’} the binary decision variable indicating whether the the stimulus orientation is clockwise or counter-clockwise of the decision boundary. The observer is assumed to solve two perceptual tasks in sequence. After stimulus presentation, the observer first performs the discrimination task using Bayesian inference with regard to a uniform loss function,

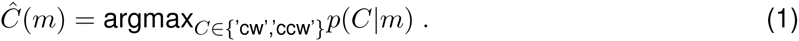

The posterior distribution *p*(*C*|*m*) is computed using Bayes’ rule, thus

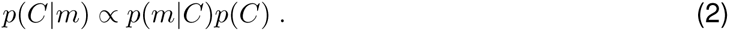

We set *p*(*C*) = 0.5 for both values of *C* since the two choices are equally likely in the Experiments 1-3 and compute *p*(*m*|*C*) by marginalizing over all stimulus orientations:

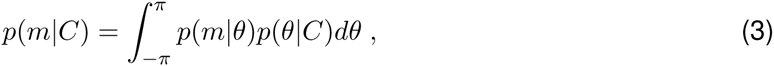

where *p*(*θ*|*C*) is the experimental distribution of stimuli for each choice. We assume the measurements *m* to be corrupted by additive Gaussian noise. Thus *p*(*m*|*θ*) is Gaussian with mean *θ* and a standard deviation *σ* that is monotonically dependent on the distribution width of the stimulus orientation array.

Second, the observer solves the subsequent estimation task by computing the mean of the posterior distribution (*i.e*. minimizing squared error), conditioned on both the sensory measurement m and their choice in the preceding discrimination task *Ĉ*:

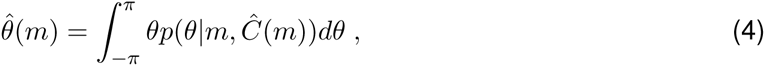

where

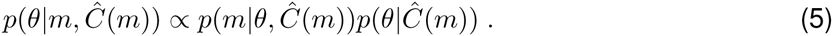

The estimation mean and the full distribution of the estimates can be computed by marginalizing Eq. (4) and Eq. (5), respectively, over the measurement distribution *p*(*m*|*θ*).

### Model fit

The conditioned Bayesian model was fit using a maximum likelihood procedure. See Supplement for more details. The predictions for the model by Jazayeri and Movshon are qualitative and not based on numerical fits.

### Code availability

Computer code (MATLAB) for the model implementation is available upon request.

---

**Extended Data Figure E1:**
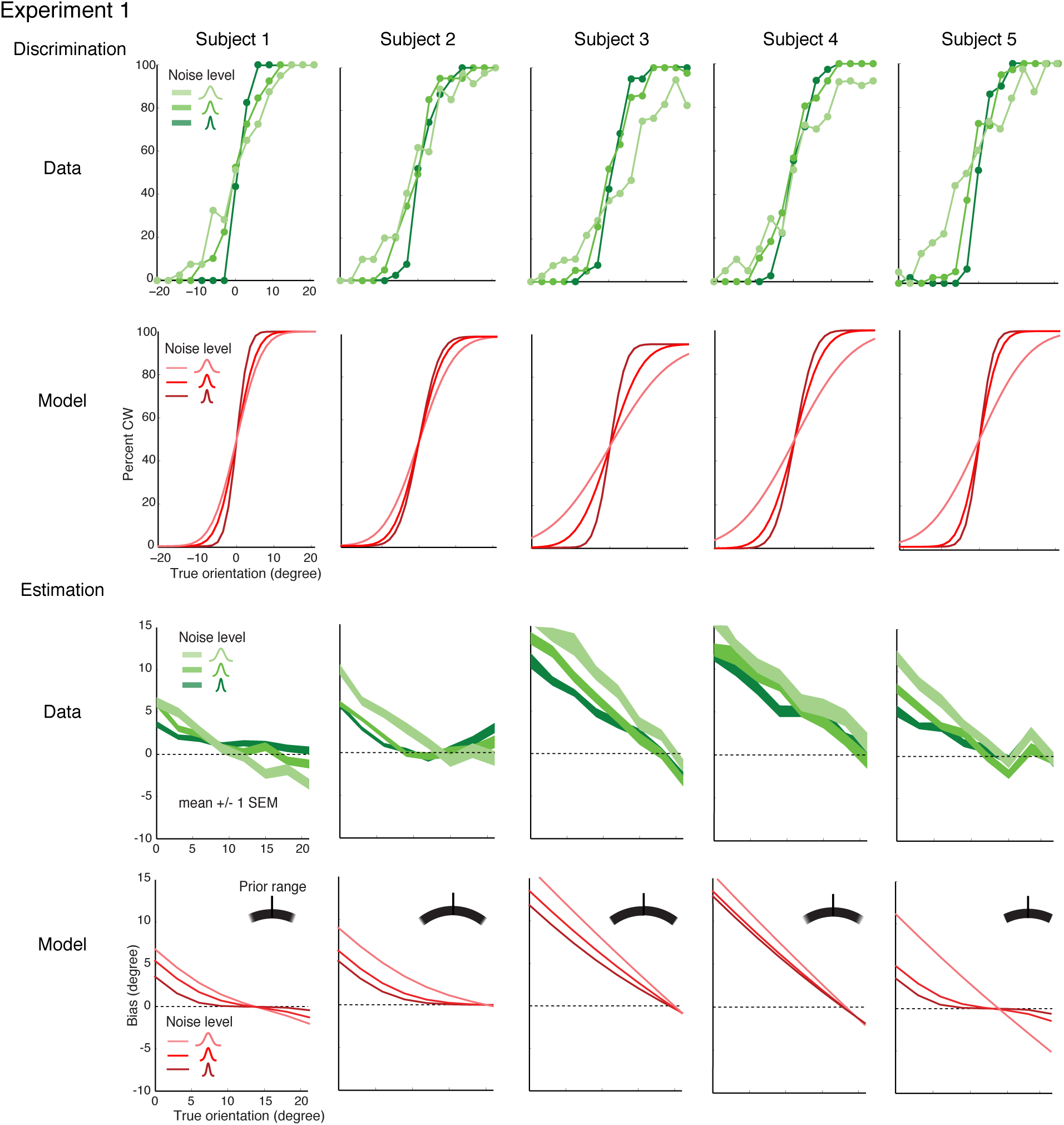
Experiment 1: Joint model fits for individual subjects. The conditioned Bayesian observer model was jointly fit to the discrimination and estimation data of individual subjects. Top two rows: Discrimination judgments of Subjects 1-5 (green points) together with the predicted psychometric curves of the model (red curves). Slopes of the psychometric curves were consistently steeper for higher stimulus noise. Subjects show large individual variations in overall noise sensitivity. Bottom two rows: Measured bias curves (correct trials only) and the corresponding model fits are shown. The bias magnitudes varies widely across subjects. However, the variations corresponds to individual differences in perceptual noise and is nicely predicted by the model. In addition, the bias curves consistently intersect around 20 degree for all subjects except the non-naïve Subject 1. This indicates that naïve subjects similarly overestimated the stimulus range as indicated by their fit stimulus priors (black arcs).

**Extended Data Figure E2:**
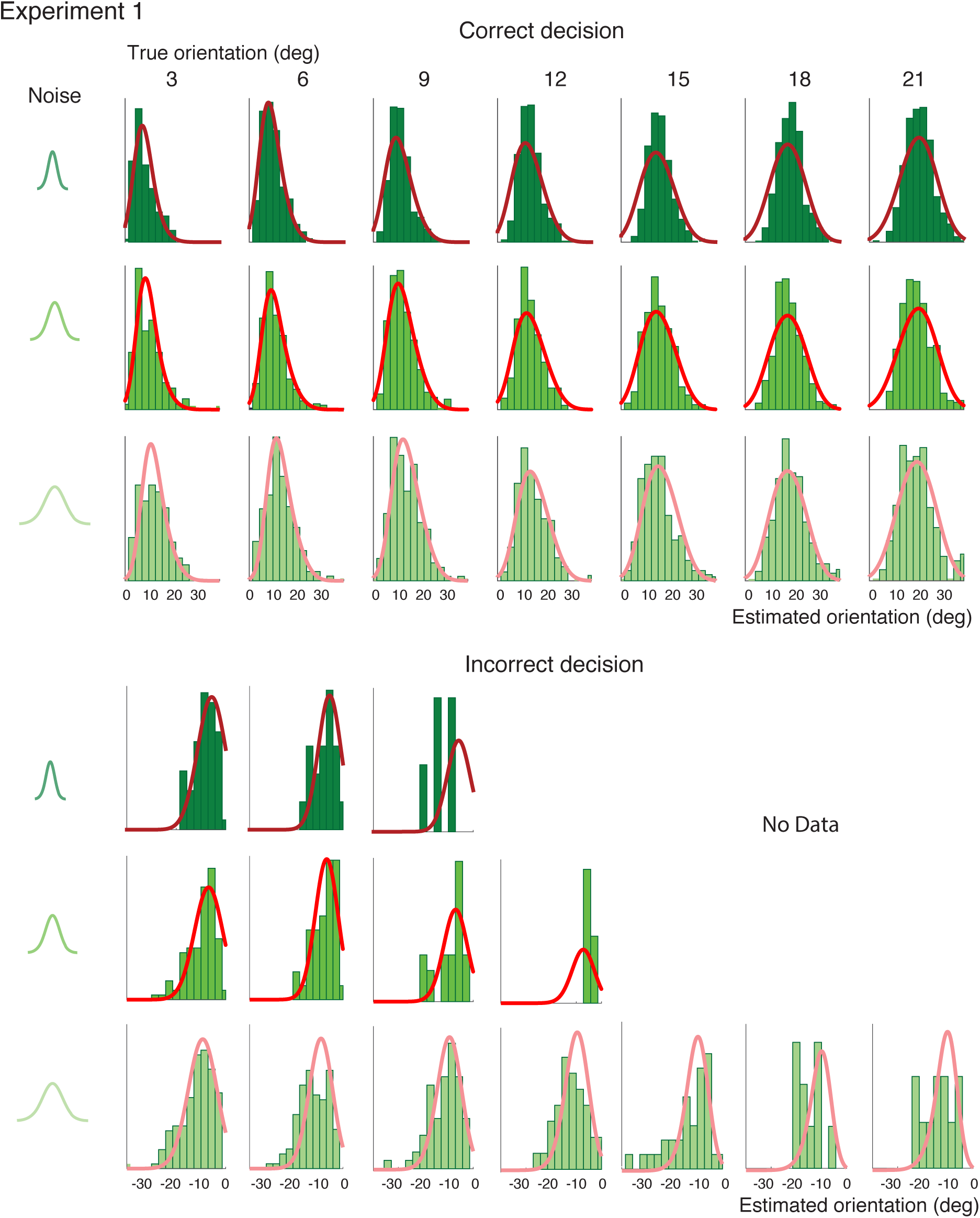
Experiment 1: Full distributions of estimates and model fits. The histograms of subjects ‘ orientation estimates at each noise level and stimulus orientation are shown for the average subject, together with the predicted distributions of the conditioned Bayesian observer model. The model closely fits the distributions for both, the correct (top) as well as the incorrect trials (bottom). Note that incorrect trials are increasingly less frequent (to completely absent at low noise) for stimulus orientations that are increasingly farther away from the discrimination boundary.

**Extended Data Figure E3:**
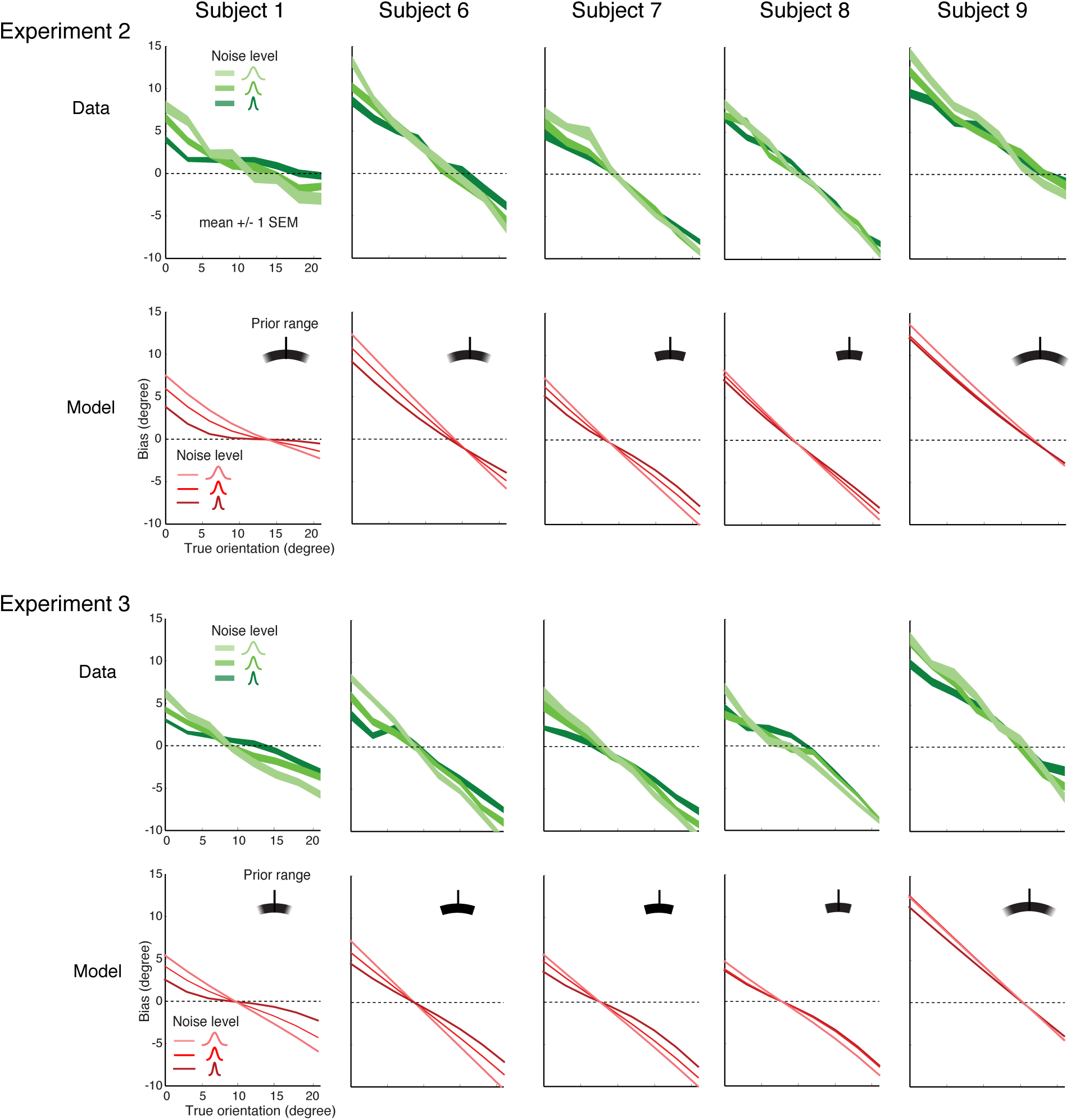
Experiment 2 and 3: Estimation data and fits for the same set of subjects. The measured bias curves in Experiment 2 are similar to those in Experiment 1 (Extended Data Figure 1) except that for most subjects the point where the bias curves intersect the x-axis (zero bias) is shifted toward the discrimination boundary. This difference is captured by the model in form of a prior (black arcs) that is narrower compared to the prior for Experiment 1 due to the explicit display of the total stimulus range in Experiment 2 (see Fig. 2a). Importantly, although the bias curves vary substantially between individual subjects, subjects show a remarkable consistency across both Experiment 2 and 3: The magnitude and shape of the bias curves are similar as well as the fit stimulus prior. Data are well fit by our conditioned Bayesian observer model.

## Supplement

### Experimental setup

#### General method

Subjects sat in a darkened room in front of a special purpose computer monitor (VIEWPixx3D, refresh rate of 120 Hz and resolution of 1920 × 1080 pixels). Viewing distance was 83.5 cm and enforced with a chin rest. We programmed all experiments in Matlab (Mathworks, Inc.) using the MGL toolbox (http://justingardner.net/mgl) for stimulus generation and presentation. The code was run on an Apple Mac Pro computer with Quad-Core Intel Xeon 2.93 GHz, 8GB RAM. Subjects were asked to fixate at the fixation dot whenever it appeared on the screen. Before subjects did the main experiments, they each had 2-3 training sessions during which they familiarized themselves with the discrimination and the estimation task. After that, each subject either completed 1800 trials in 3-4 sessions for Experiment 1 or completed 3600 trials in 6-8 sessions for Experiment 2 and 3. This is equivalent to 40 trials per each of the 15 stimulus orientations and the three noise conditions. Each session lasted approximately 50 minutes.

#### Experiment 1

Five subjects (Subjects 1-5) participated in Experiment 1. In each trial, subjects viewed a white fixation dot (diameter: 0.3^*o*^) and two black marks (length: 3^*o*^, distance from fixation: 3.5^*o*^) indicating a decision boundary whose orientation was randomly chosen around the circle. After 1300 ms, an array of white line segments (length: 0.6^*o*^) was presented for 500 ms. The array consisted of two concentric circles centered on the fixation: the outer (diameter: 3.8^*o*^) contained 16 line segments and the inner (diameter: 1.8^*o*^) contained 8 line segments. A random jitter (from −0.15^*o*^ to 0.15^*o*^) was independently added to the x-y coordinates of each line segment. The orientation of each line segment was drawn from a Gaussian distribution with mean given as one of 15 stimulus orientations relative to the boundary (from −21^*o*^ to 21^*o*^ in steps of 3^*o*^) and standard deviation as one of 3 values ([0^*o*^,6^*o*^ and 18^*o*^]). After the stimulus disappeared, subjects were asked to indicate whether the average orientation of the array was clockwise or counter-clockwise relative to the boundary by pressing a corresponding button. If subjects did not respond within 4 seconds, the current trial was skipped. If they responded within 4 seconds, subjects then subsequently were tasked to indicated their perceived average orientation of the array by adjusting a reference line with an analog stick of a gamepad (PS4 DualShock 4). Each trial was followed by a randomly chosen inter-trial interval of 300 ms to 600 ms duration (blank screen).

#### Experiment 2

Five subjects (Subject 1 and Subjects 6-9) participated in Experiment 2. The procedure was identical to Experiment 1 except that at the beginning of each trial, a prior cue consisting of a gray arc was presented for 800 ms. The arc (width: 0.2^*o*^, eccentricity from fixation: 3.5^*o*^)always spanned the range −21^*o*^ to 21^*o*^ relative to the discrimination boundary. Subjects were instructed that the stimulus orientation was guaranteed to occur anywhere within this range with equal probabilities.

#### Experiment 3

The same five subjects in Experiment 2 also participated in Experiment 3. The procedure was identical to Experiment 2 except for the following: First, the full prior cue (gray arc) was present only for 500 ms, after which it was reduced to a colored arc that only spanned the orientation range at the side of the discrimination boundary where the stimulus orientation in this trial would occur. This colored cue indicating the correct answer (’cw’ or ‘ccw’) was shown for 300ms. The color (red or green) was randomly assigned and uncorrelated with the stimulus orientation. Second, instead of the orientation discrimination task, subjects were tasked to recall the color of the answer cue.

### Fitting procedure

We jointly fit the model to the data of both the decision and estimation task by maximizing the likelihood of the model given the data.

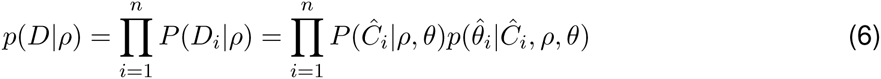

where *D* is the data, *ρ* is the parameters of the model, *θ* is the true orientations, *Ĉ*_*i*_ is the decision outcome, *θ̂*_*i*_, is the orientation estimate, *i* is the trial index and *n* is the number of trials. Thereby, the probability of the decision *P*(*Ĉ*|*θ*), and the distribution of estimates conditioned on the decision *p*(*θ̂*|*Ĉ*, *θ*) are computed by marginalizing Eqs. (1) and (5) over the measurement distributions *p*(*m*|*θ*) (Gaussian).

Our model contains a total of 8 parameters including:

- standard deviations for the 3 noise levels of the stimuli (additive Gaussian noise)
- lapse rate and guess rate for the discrimination task
- width and smoothness of the prior distribution over orientation
- standard deviation for memory noise (additive Gaussian)

The Nelder-Mead simplex algorithm is used to minimize the term −*log*(*p*(*D*|*ρ*)). Twenty iterations of the optimization procedure were performed using randomized initial parameter values in oder to obtain the best fitting model.

